# Elucidating vimentin interaction with zinc ions and its interplay with oxidative modifications through crosslinking assays and molecular dynamics simulations

**DOI:** 10.1101/2021.02.12.430929

**Authors:** Andreia Mónico, Joan Guzmán-Caldentey, María A. Pajares, Sonsoles Martín-Santamaría, Dolores Pérez-Sala

## Abstract

The intermediate filament protein vimentin is involved in essential cellular processes, including cell division and stress responses. Vimentin oxidative modifications impact network reorganization and its single cysteine residue, Cys328, acts as a redox sensor. Vimentin binds zinc, which influences its assembly by undefined mechanisms. Here, results from combined biochemical and molecular dynamics studies support that zinc ions interact with Cys328 in its thiolate form, whereas Glu329 and Asp331 stabilize zinc coordination. Vimentin oxidation can induce disulfide crosslinking, implying a close proximity of cysteine residues in certain vimentin associations, validated by our computational models. Notably, micromolar zinc concentrations selectively prevent Cys328 alkylation and crosslinking. These effects are not mimicked by magnesium, consistent with the fewer magnesium ions hosted at the cysteine region. Altogether, our results pinpoint the region surrounding Cys328, highly conserved in type III intermediate filaments, as a hot spot for zinc binding, which modulates Cys328 reactivity and vimentin assembly.

## Introduction

Vimentin is a type III intermediate filament protein that forms robust filaments in mesenchymal cells. The intermediate filament network maintains an intimate crosstalk with the other cytoskeletal systems (microfilaments and microtubules) and modulates essential cell properties, from cell size to migration, division and elasticity ^1–3^. Importantly, the vimentin network finely responds to electrophilic and oxidative stress, a function in which its single cysteine residue (Cys328) plays an important role ^4, 5^. In addition, vimentin is involved in disease by various mechanisms, including acting as a cellular receptor for pathogens, such as bacteria and viruses ^6^, regulating the immune response ^7^ or playing a key role in epithelial-mesenchymal transition during tumorigenesis ^8^.

The cellular organization and architecture of vimentin filaments are still not completely understood. In contrast to the polarity observed in microfilaments and microtubules, vimentin filaments are non-polar and can grow at both ends. Moreover, they are highly dynamic in cells and can exchange subunits along their length and undergo severing and reannealing ^9, 10^. Nevertheless, the regulation and/or cofactors modulating these processes need further characterization.

Vimentin polymerization in vitro has been exhaustively analyzed by a variety of techniques, including electron microscopy (EM) and cryo-EM, atomic force microscopy, small angle X-ray scattering, sedimentation velocity and total internal reflection fluorescence ^11–16^. A consensus regarding the assembly pathway has emerged according to which, parallel vimentin homodimers engage in a staggered manner in antiparallel tetramers, eight of which laterally associate to give a unit-length filament (ULF) (Figure 1A). These ULFs assemble head to tail to form filaments. During or after this elongation process, filaments “mature”, undergoing an internal rearrangement by which the initial ULF diameter (~16 nm) diminishes down to 10-12 nm, considered the diameter of normal filaments ^11, 17^.

**Figure 1:**
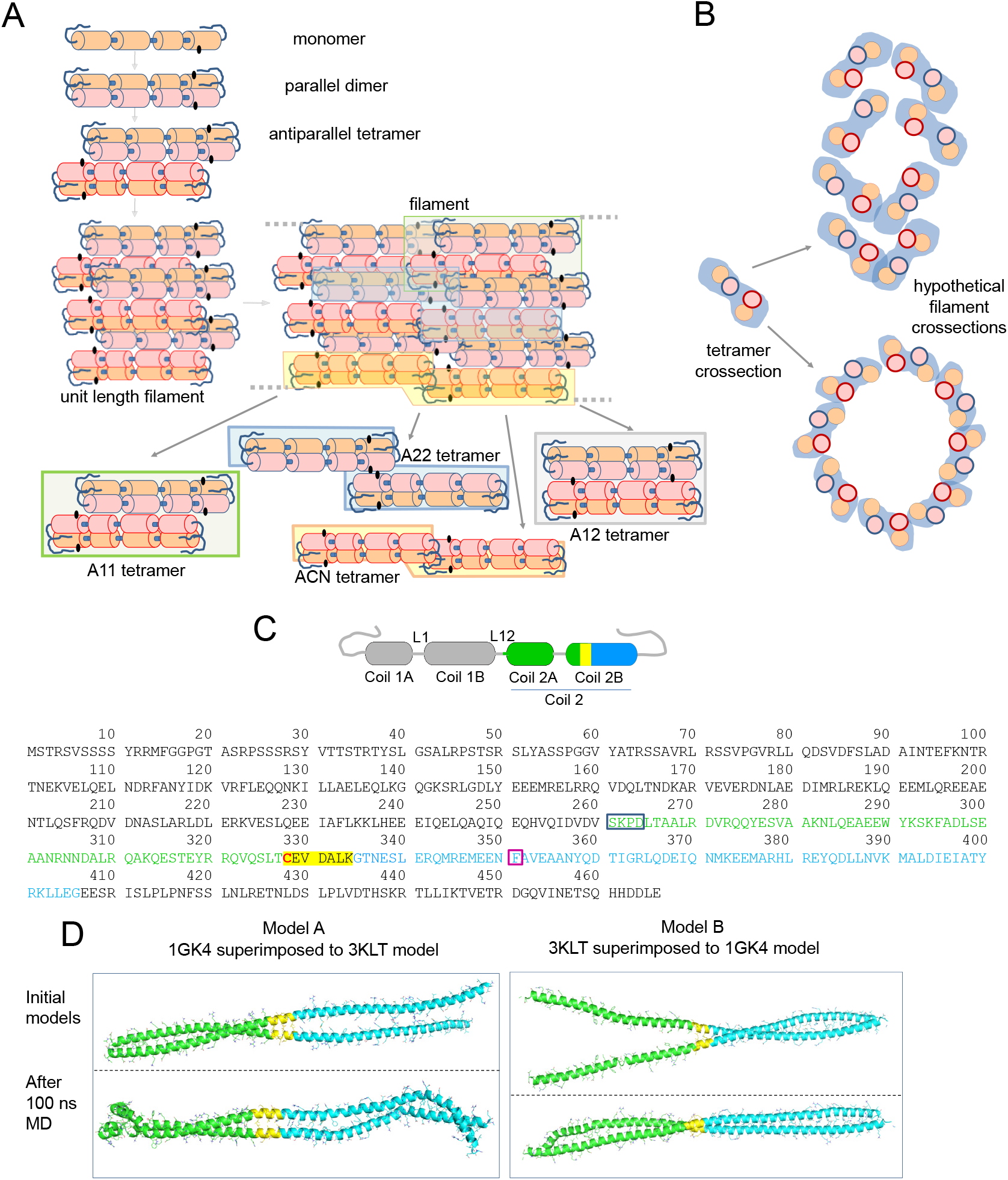
(A) Scheme of vimentin assembly and the proposed configurations of vimentin tetramers in filaments (please see text for details). A cartoon view interpretation of the assembly of two unit length filaments (ULF) is presented (labeled as “filament”) showing the potentially key position of the cysteine residue (black dots), which could fall close to the region involved in the connection of two ULFs. Within assembled filaments, vimentin dimers can coincide in different relative positions, thus resulting in several tetrameric configurations, some of which are outlined in the figure by color shadowing and extracted below for easier visualization. (B) Cartoon of a tetramer transversal view and hypothetical dispositions of tetramers in a transversal section of a filament. Please note that these schemes are inspired in previous works ^14, 93^ and are not based on molecular dynamics simulations. (C) Scheme of a vimentin monomer showing the distribution in coiled coil segments and linkers according to ^71^. The segments used for building the models shown in (D) are highlighted by a color code: amino acids present in the 3KLT and 1GK4 crystal structures are represented in green and blue, respectively; the region shared between both crystal structures is highlighted in yellow and the cysteine residue is shown in red. The position of those segments in the full vimentin sequence is depicted underneath by the same color code. In addition, color boxes outline the amino acids integrating the linker (blue) and the position of the stutter (purple). (D) Representation of both models (A and B) built from the superimposition of 3KLT (green) and 1GK4 (blue) crystal structures on the shared region of both crystal structures (yellow). The upper pictures represent the initial model and lower pictures the model after 100 ns of MD simulation.

Vimentin monomers consist of three domains, head, rod and tail, of which, the rod is mainly alpha-helical, whereas the head and tail domains lack a defined structure. Being a coiled-coil protein for a high proportion of its sequence, solving the crystal structure of vimentin poses great difficulties, for which it has been addressed through a “divide and conquer” approach. In these studies, the structure of some segments in different degrees of association, i.e., dimers or tetramers, has been solved (please, see ^18^ for review). In addition, elegant early works using amino-reactive crosslinkers, like disuccinimidyl tartrate (DST), have yielded insight into the disposition of vimentin dimers in tetramers and of tetramers within filaments ^19, 20^. The results obtained suggested the predominance of an arrangement involving the amino-terminal overlap of the two dimers, the so-called A11 mode, in preparations of soluble vimentin, and an increased abundance of other alignment modes coexisting in vimentin filaments, namely, a C-terminal overlap, the A22 mode, a head to tail overlap, the ACN mode, or a complete antiparallel overlap, the A12 mode. The way these associations could be present within filaments is illustrated in a cartoon view of a two-ULF assembly (Figure 1A). More recently, deuterium exchange approaches have yielded information on the vimentin head domain ^21^, whereas electron paramagnetic resonance has been used to obtain insight into the structure of the head ^22^ and tail domains ^23^. Information collected through these approaches has been integrated with molecular modeling studies providing our current view of vimentin assembly ^14, 24^ (reviewed in ^25^). Moreover, information from various intermediate filament proteins has contributed to the understanding of filament structure ^26, 27^. Most available models of the vimentin tetramer adopt the A11 configuration, in which an anchoring knob–hydrophobic pocket mechanism involving the 1B segment, identified in several intermediate filament proteins including vimentin, provides stability ^28^ (see ^27^ for review). In turn, in ULF or filaments, a parallel stacking of tetramers is proposed, usually represented as a tubular structure ^14, 29, 30^ (Figure 1B), although their exact disposition and relative positions with respect to the longitudinal axis, are not understood.

Vimentin contains a single cysteine residue, Cys328, located in coil 2B (Figure 1C), which constitutes a hot spot for modification by oxidants and electrophiles and is required for oxidative vimentin filament disruption in vitro and network rearrangement in cells, thus acting as a redox sensor ^4, 31, 32^. However, the precise location of Cys328 residues within filaments and how their modifications impact filament organization remains unknown. Crystal structures of segments of coil 2B including Cys328 (residues 254-411), show the lateral chain of Cys328 facing outwards the dimer ^33, 34^, a fact that is also observed in models available through databases (e.g. P02543) ^29, 30^. Some of these crystal structures have yielded tetramers of the peptides used, which do not correspond with any of the assemblies identified in crosslinking studies using full length vimentin, and therefore represent “non-physiological organizations” ^34^. Moreover, in some cases, the length of the segment solved precludes the complete identification of interactions involving Cys328. Additional data on coil 2B interactions obtained by site-directed spin labeling of residues around Cys328 and electron paramagnetic resonance ^35^, reveal residue 348 as the strongest point of interaction and overlap (within 1.6 nm) between two antiparallel dimers. However, vimentin, as well as other intermediate filament proteins, has been reported to undergo oxidative crosslinking through its single cysteine residue ^36^. The major product of this crosslinking is a disulfide-bonded oligomer, likely arising from the interaction between adjacent native tetramers, which migrates as a dimer in non-reducing SDS-PAGE ^5, 36^. This suggests that in certain conformations of soluble or filamentous vimentin, cysteine residues, likely from different tetramers, are located at a distance amenable to disulfide crosslinking. Nevertheless, the correspondence of these conformations with the previously described modes of alignment is not straight forward.

The vimentin molecule behaves as a polyelectrolyte due to the abundance of negatively charged residues. This feature promotes its interaction with divalent cations, such as calcium and magnesium, which, at millimolar levels, influence vimentin in vitro assembly and have been reported to behave as crosslinkers for certain regions of the protein ^37, 38^, affecting the properties of the network ^39^. In recent works, we observed that zinc interacts with vimentin at micromolar concentrations ^4^, modulating vimentin in vitro assembly in an ample range of concentrations ^40^. Moreover, zinc levels influence the organization of the network and its resistance to oxidants in cells ^4^. Thus, we proposed that zinc could interact with vimentin at several sites, of which, the region surrounding Cys328 could be especially relevant ^41^. Zinc(II) ions interact with proteins playing catalytic, regulatory or structural roles, usually coordinating with amino acids such as cysteine, histidine or carboxylic amino acids ^42, 43^. Moreover, the interaction of Zn(II) ions with cysteine residues can modulate their reactivity and impact their redox regulation ^44^. Therefore, given the importance of Cys328 in vimentin assembly and response to oxidative stress, elucidating its interaction with zinc will contribute to the understanding of vimentin regulation.

Here, we propose a computational model of the pivotal sequence of vimentin comprised between Asp264 and Gly406, in order to gain insight into the organization of the protein and its interaction with zinc. Moreover, we have experimentally addressed the interplay of bifunctional cysteine crosslinkers and zinc. Our results provide a hypothesis for the organization of the vimentin central domain, its putative interaction with zinc and its involvement in the formation of dimers of dimers under oxidative stress conditions, in which the cysteine residues could be spatially close.

## Results and Discussion

### Computational modeling of a vimentin dimer and its interaction with zinc

Zinc binding sites in proteins often involve cysteine residues or carboxylic amino acids ^45, 46^. Based on our previous work ^4^, we focused our attention into the Asp264-Gly406 region of vimentin surrounding the cysteine residue in order to explore its potential interaction with zinc (Figure 1C). Two different models of vimentin dimer, namely models A and B, were computationally built from the X-ray structures of PDBs 3KLT ^34^ and 1GK4 (Figure 1D, see Materials and Methods for details), and submitted to 100 ns molecular dynamics (MD) simulation (Figures 1D and S1). Interestingly, both models adopted a typical coiled-coil configuration during the simulation time. However, the final structure of model A cannot be considered as a coiled-coil, since both ends of the protein came into contact. In contrast, model B achieved the desired coiled-coil structure after the 100 ns simulation (Figures 1D), and remained stable during the whole simulation time when it was extended until 200 ns. Interestingly, the dimerization interface between monomers in this model is mostly composed of hydrophobic residues, similarly to other related dimers, as that of vimentin rod 1B (PDB 3UF1) ^47^. Of note, in this simulation, the stutter, located at residue 351, in which the coiled motif has been reported to be disrupted, adopts a parallel distribution, not a coiled coil motif. This is consistent with previous observations ^33^. Moreover, the region corresponding to PDB 3KLT is maintained very similar to the X-ray structure reported, without showing relevant structural changes.

Therefore, model B was used to interrogate the hypothesis of a potential interaction of zinc atoms in this region (Figure 2A). Firstly, mapping the electrostatic potential surface of this dimer allowed the location of 23 possible coordination sites (hot spots) for zinc ions, i.e. regions with predominant negative density charge (Figure 2A). To saturate the system, we introduced additional ions (up to 50) around the vimentin dimer. Zinc ions were manually relocated in order to fill all the binding hot spots and the systems submitted to 30 ns of MD simulation. Thirteen of the 50 zinc atoms remained in close contact with vimentin residues for more than 90% of the simulation time, indicating a good stability of zinc-vimentin complexes (Figure S2). All of these zinc ions were interacting with side chains from Asp or Glu residues exposed to the solvent. Of note, in some cases, zinc atoms could interact with more than one residue located either on the same chain or on different chains (see below).

**Figure 2:**
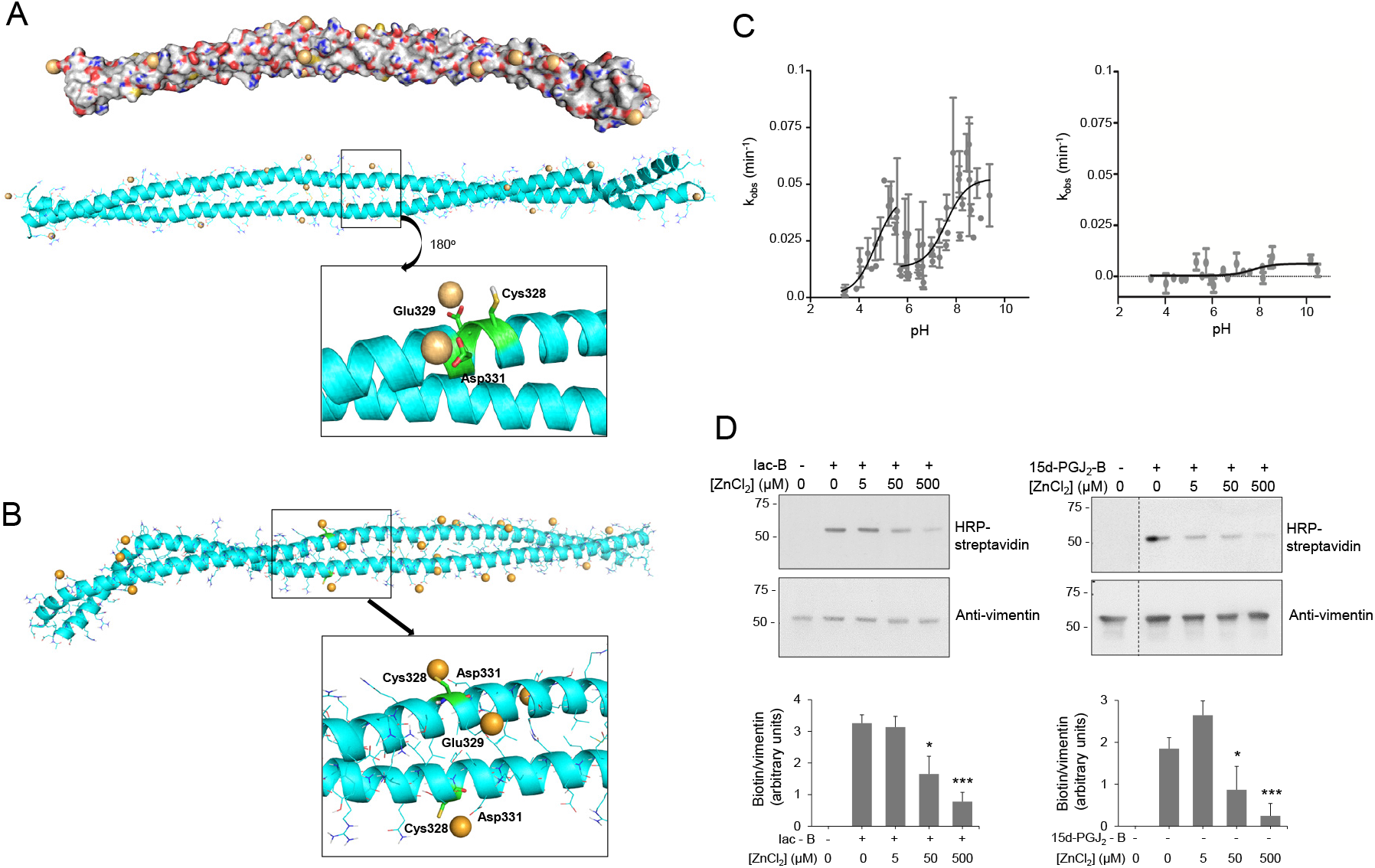
Vimentin dimer and its interaction with zinc. (A) Zinc binding at vimentin B dimer. Filled and ribbon views of dimer B are shown. After 100 ns MD simulation, zinc atoms, shown as orange spheres, were placed at electronegative regions. Below, the region near Cys328 is displayed in detail illustrating the interaction of zinc atoms with Glu329 and Asp331. After 30 ns of MD simulation, two Zn atoms remained close to Cys328, interacting with Glu329 and Asp331 during the whole simulation time. (B) Zinc binding at vimentin B dimer, with Cys328 in its thiolate form. In this case, the cysteine interacts directly with zinc atoms as displayed in the detailed view. (C) Experimental determination of vimentin Cys328 pKa. Kinetics of MBB modification of vimentin wt (left panel) and Cys328Ser mutant (right panel) were measured at different pHs in triplicate using a Varioskan plate reader and the k_obs_ calculated for each case. The pKa was obtained from the representation of k_obs_ values against the final pH of the reaction. (D) Protection of vimentin modification by zinc. Purified vimentin wt (4.3 μM final concentration) was incubated with ZnCl_2_ at the indicated concentrations for 1 h at r.t., before adding vehicle, biotinylated iodoacetamide (10 μM, Iac-B, left panel) or biotinylated 15d-PGJ_2_ (1 μM, 15d-PGJ_2_-B, right panel), and incubating an additional hour at 37°C. Incubation mixtures were run on SDS-PAGE gels, electro-blotted and incorporation of the biotinylated compounds assessed by biotin detection with HRP-conjugated streptavidin. Vimentin was detected by western blot. Dotted lines indicate where lanes from the same gel have been cropped. The position of molecular weight standards (in kDa) is indicated on the left. The ratio of the biotin and the vimentin signals is displayed in the graph as mean values ± SEM of four independent assays. *p <0.05 vs Iac-B or 15d-PGJ_2_-B at 0 μM ZnCl_2_; *** p < 0.0005 vs Iac-B or 15d-PGJ_2_-B at 0 μM ZnCl_2_ by Student’s *t*-test.

Other proteins have been found to interact with multiple zinc atoms, mainly through carboxylic residues, which can play a structural role stabilizing subunit interactions ^48^. Interestingly, two of the zinc ions bound to the Asp264-Gly406 region of vimentin were found to establish stable interactions with Glu329 and Asp331, which are located next to Cys328 (Figure 2A, inset). These stable interactions may pinpoint an important zinc coordination site that could contribute to Cys328 protection against modification by alkylating agents ^4^.

Importantly, cysteine residues can exist either in thiol (−SH) or thiolate (−S^−^) forms depending on their pKa and the environmental pH. In our initial model B, we considered the neutral form of Cys328 thiol group (Figure 2A). Therefore, we next modeled vimentin with the cysteine residue in its thiolate form (Figures 2B and S3). MD simulations were performed following the same protocol as for vimentin with Cys328 in its thiol form (-SH). Interestingly, almost the same number of zinc ions remained close to the vimentin dimer along the simulation time. Also two zinc atoms were located at the hot spots involving Glu329 and Asp331; however, in this case, the Cys328 thiolate entered into the zinc coordination shell maintaining more stable interactions (length of the Zn-S coordination bond 2.5 Å, Figures 2B and S3). These observations point to the complexity of the vimentin-zinc interaction. On one hand, multiple vimentin species may coexist depending on the oligomeric state of the protein and on the ionization status of the cysteine residue. On the other hand, given the importance of Cys328 in vimentin assembly and redox sensing ^4, 5^, the interaction could have implications in the reactivity and/or accessibility of this residue, thus affecting its oxidation or modification by electrophiles.

### Determination of Cys328 pKa

Vimentin Cys328 has been shown to be the target of several posttranslational modifications, including glutathionylation, lipoxidation and nitrosylation ^5, 49, 50^, which typically affect cysteine residues with a low pKa, and therefore are present in their thiolate form in substantial proportions at physiological pH. The thiolate form is more reactive than the thiol and therefore, cysteine residues with low pKa are considered to be particularly reactive with electrophilic compounds and/or more prone to oxidation. In order to assess the relative abundance of the thiol and thiolate forms of Cys328 under conditions close to physiological pH, we set out to determine its pKa. The computational calculation yielded pKa values of 9.3 for cysteines in the dimer, whereas for the dimer of dimers (see below), values of 9.3 and of 10.0 were obtained for cysteines facing outwards and the interdimer space, respectively (Figure S4). These values are in accordance to those recently obtained for Cys328 in the context of the 327-329 or the 321-334 vimentin peptides, for which pKa values of 8.5 ± 2.2 were obtained following different computational approaches ^51^. Since theoretical and experimental pKa values often differ ^52^, we undertook the experimental determination of Cys328 pKa (Figure 2C). For this purpose, we set up a procedure based on the modification of cysteine residues by the fluorescent compound monobromobimane (MBB) ^53^. Incubation of soluble vimentin with MBB led to the pH-dependent incorporation of the fluorescent probe, as shown upon calculation of k_obs_ values for the reactions carried out at each pH (Figure 2C). Interestingly, the results obtained were consistent with the existence of two ionization steps for Cys328 in soluble vimentin, rendering pKa values of 4.61 ± 0.36 and 7.39 ± 0.41 (mean ± SD) (Figure 2C, left). Therefore, the experimental pKa values are much lower than the theoretical values obtained from the model or from cysteine in solution, and indicate that a substantial proportion of Cys328 exists in its thiolate form at physiological pH. Incorporation of MBB into Cys328 is specific, as a Cys328Ser mutant, used as a control, showed only background modification and nearly null pH dependence, evidenced by the dispersion of the data and the low R^2^ of the curve fit (R^2^ = 0.07249) (Figure 2C, right). The pKa of cysteine residues is greatly influenced by the chemical environment, including the proximity of charged residues or hydrogen bonding ^52^. Cysteines with particularly low pKa values have been reported at the active sites of the thioredoxin superfamily ^54^ or of cysteine proteases ^55^. However, the current available vimentin structures do not provide a complete view of the environment surrounding the lateral chain of Cys328 to discern the cause of the low pKa values detected. Moreover, the observation of two different pKa values for vimentin could be related to the presence of different oligomeric or conformational species in the protein preparations. It is well known that soluble vimentin can exist in several oligomeric species, which may include tetramers and octamers ^11, 20, 56^. Therefore, our results could be interpreted on the basis of the coexistence of a variety of oligomeric or conformational species in soluble vimentin preparations, in which the environment of Cys328 could be significantly different, affecting its pKa. In order to strengthen these observations, the alkylation of vimentin by biotinylated iodoacetamide (Iac-B) was assessed at pH ranges around the pKa values determined by MBB using a gel-based assay (Figure S4). This semiquantitative assay confirmed the existence of two pH ranges for Cys328 modification, with more intense Iac-B incorporation for thiolates with pKa 7.39.

Taken together these results indicate that the pKa of Cys328 falls within the physiological pH range, suggesting that a significant proportion of cysteine residues in vimentin molecules will occur in thiolate form in cells and supporting the behavior of this residue as a hot spot for oxidative modification and therefore, redox sensing ^4, 57^.

### Zinc protects vimentin from cysteine alkylation and lipoxidation

To substantiate the importance of the interaction of zinc with the Cys328 region, we assessed the extent of modification by Iac-B after preincubating the protein with ZnCl_2_ (Figure 2D). Micromolar concentrations of ZnCl_2_ prevented subsequent vimentin alkylation by Iac-B in a concentration-dependent manner (Figure 2D, left panel). Furthermore, ZnCl_2_ protected vimentin from lipoxidation by the electrophilic prostaglandin 15d-PGJ_2_ (Figure 2D, right panel), which selectively modifies Cys328 ^5^. Protection of cysteine residues from alkylation has been previously proposed as an indication of zinc binding ^58^. Thus, these results support the interaction of zinc with vimentin in the proximity of the cysteine residue. Nevertheless, the possibility that zinc could induce vimentin aggregation and/or conformational changes reducing the accessibility of Cys328 to modification should also be considered ^40^. In contrast, we have previously reported that NaCl-induced polymerization does not preclude vimentin oxidation or lipoxidation, which indicates that the protection is not merely due to lower cysteine accessibility in oligomeric structures, and suggests a distinct role for zinc ^5^. Interestingly, ZnCl_2_ concentrations providing protection against Cys328 modifications in our assays are in the order of the total concentrations found in plasma ^59^ or in cells ^60^, i.e., approximately 12 and 200 μM, respectively. Nevertheless, most of zinc is bound to proteins and its levels vary greatly depending on the subcellular compartment. Thus, picomolar to nanomolar concentrations of free zinc have been measured in the cytoplasm, whereas in certain organelles such as lysosomes or insulin-storing granules, zinc accumulates reaching concentrations between 1 and 100 μM ^61, 62^. Therefore, the modulation of vimentin modifications by zinc in the cell may change notably depending on the local ion concentrations. From the chemical point of view, a complex interplay may exist between zinc binding and cysteine reactivity. On one hand, zinc binding can lower the pKa of cysteine residues stabilizing the negatively charged thiolate anions and making them more reactive towards oxidants and electrophiles ^43, 63^. In any case, a low pKa implies a higher fraction of reactive thiolates, but it also may imply a lower nucleophilicity ^64^. Therefore, the impact of zinc binding on thiol accessibility and reactivity needs to be evaluated on an individual basis. Indeed, there are examples of protective effects of zinc binding on cysteine modification, e.g., alkylation by iodoacetamide or oxidation ^58, 65^, but also cases in which modification is promoted, e.g., persulfidation by H_2_S ^65^. This interplay can be even more complex in the cellular context, where prooxidant and antioxidant actions of zinc can take place depending on the cellular context and zinc levels ^44^.

### Modeling of a vimentin Cys328Ser mutant

Mutation of cysteine to serine in proteins is a widely used strategy to assess the importance of thiol moieties in protein function and signaling. Regarding vimentin, we have previously shown that a Cys328Ser mutant is competent for filament formation in vitro and confers resistance to the alterations of vimentin filaments induced by electrophilic agents ^5^, although it shows subtle but significant differences in polymerization features and response to zinc supplementation compared to vimentin wt ^40^. Therefore, it was of interest to model a potential interaction of this mutant with zinc. The starting geometry of vimentin dimer B was used to computationally mutate Cys328 to serine (Figure 3). The resulting structure was submitted to 50 ns of MD simulation, and no major differences with respect to the wt overall structure were observed (Figure S5). However, detailed observation of the Ser328 region showed that the H-bond linking the Ser328 OH group and the CO group from the Gln324 backbone was weaker than the H-bond established between the SH and CO groups in the wild type (wt) protein, as deduced from monitoring the H-bond distance. Additionally, the electrostatic potential of the surface of the Cys328Ser dimer B was slightly more electronegative than that of the wt dimer. The main differences affected the zinc binding sites. Whereas the wt was shown above to bind two zinc ions, four zinc ions remained bound at the vicinity of the serine residues (two ions per residue) in the simulation of the mutant protein. Therefore, the vimentin Cys328Ser mutation confers resistance to disruption by oxidants ^4, 5^, and may also result in differences in the response to zinc availability.

**Figure 3.**
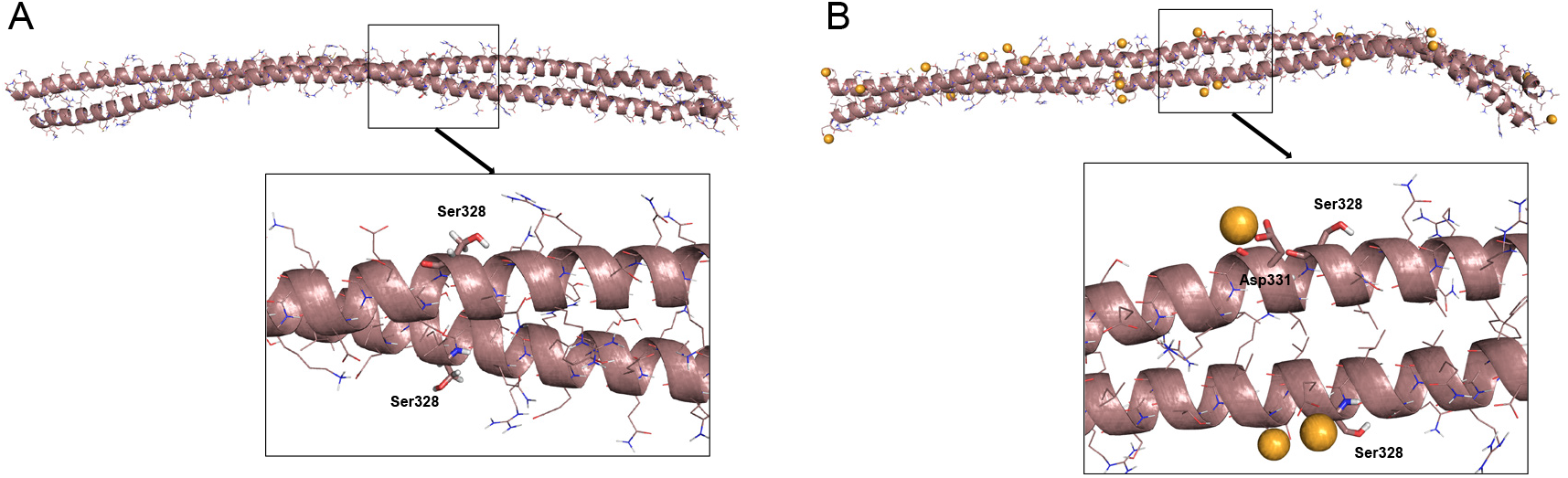
B dimer of a vimentin Cys328Ser mutant and its interaction with zinc. A) Final snapshot of a 100 ns MD simulation of the Cys328Ser mutant in water. B) Final snapshot of 100 ns MD simulation of this mutant in the presence of zinc. In both cases, lower panels show an amplified view of the region around Ser328. Zinc atoms are represented as orange spheres.

### Modeling of a vimentin dimer of dimers and its interaction with zinc

The relative position of vimentin tetramers within filaments is a complex problem not completely understood yet. Importantly, oxidative conditions have been reported to induce the formation of oligomeric species of vimentin by disulfide bonding ^5, 36^, which correlates with altered assembly. Indeed, as shown in Figure 4A, incubation of vimentin with various oxidants or cysteine-crosslinking agents, such as diamide, dibromobimane (DBB) or hydrogen peroxide, gave rise to oligomeric species migrating slightly above the 150 kDa molecular weight marker. This electrophoretical mobility, although lower than the theoretically expected, is compatible with the formation of a vimentin dimer, as previously reported ^5, 36^. Of the various agents used, diamide and H_2_O_2_ induce disulfide bonds, and therefore, reversible dimerization ^5^, whereas DBB induces irreversible crosslinking ^66^. Formation of the vimentin dimer requires the presence of the cysteine residue, as illustrated by the lack of DBB crosslinking in the Cys328Ser mutant (Figure 4B). Consistent with previous results ^4^, preincubation of vimentin with NaCl under conditions leading to polymerization into filaments resulted in an increase in the formation DBB-induced oligomeric species (Figure 4C). Taken together, these results suggest the existence of certain conformations in soluble vimentin and within filaments in which vimentin cysteines, likely from different tetramers, are located at a distance amenable to disulfide formation or DBB crosslinking. Importantly, cysteine-crosslinked oligomers also occur in other type III intermediate filament proteins, like glial fibrillary acidic protein (GFAP), which is highly homologous to vimentin ^31^. Moreover, cysteine-crosslinked heterooligomers of vimentin and GFAP or desmin have been detected in several experimental systems, both in vitro and in cells, which supports the proximity of the cysteine residues in pathophysiological cytoskeletal arrangements ^31, 67, 68^. Among the potential tetrameric associations, the A22 overlap mode would allow the shortest distance between cysteine residues in the adjacent dimers, although still not ideal for disulfide formation ^36^. However, as pointed out in ^36^, in this conformation, the stutter falls between the two cysteine residues and may impose alterations in the expected configuration in this region. Indeed, the stutter has been suggested to lead to a softer structure with less resistance to unfolding ^69^.

**Figure 4.**
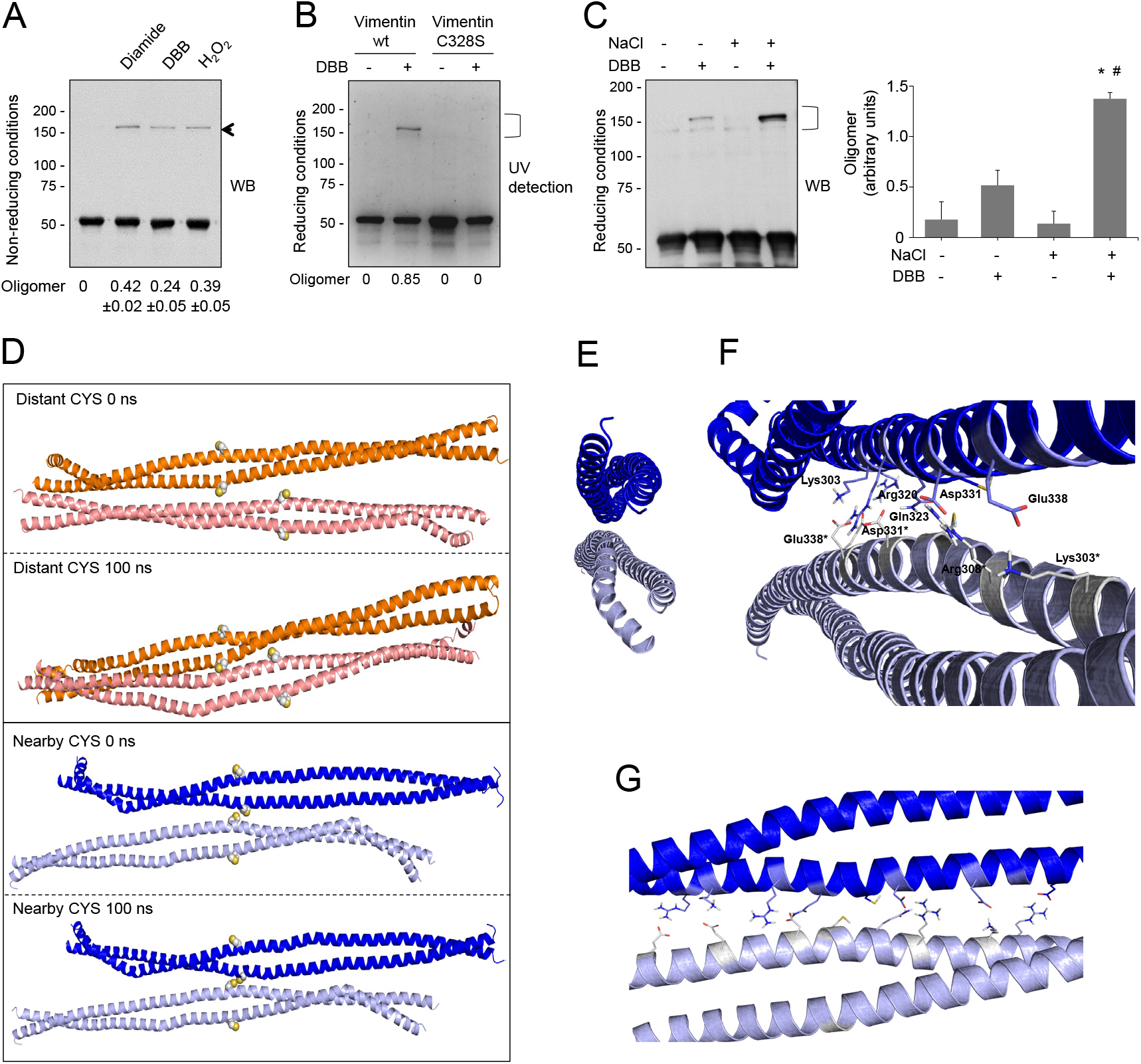
Cysteine-crosslinking of vimentin and potential conformations of the resulting dimers of dimers obtained by molecular dynamics. (A) Cysteine-crosslinking of vimentin (4.7 μM final concentration) upon incubation with vehicle, 1 mM diamide, 24 μM DBB or 1 mM H_2_O_2_ for 1 h at r.t. Incubation mixtures were analyzed SDS-PAGE under non-reducing conditions and western blot (WB). The position of the oligomer detected under these conditions is marked by an arrow. The oligomer/monomer ratios are displayed below the plot as average values ± SEM of three independent experiments. (B) Cys328 is required for DBB crosslinking. Vimentin wt or Cys328Ser (4.5 μM final concentration) were incubated with 50 μM DBB for 1 h at r.t, and subjected to SDS-PAGE, followed by Sypro-Ruby staining for detection of vimentin monomeric and oligomeric (marked by a bracket) species under UV light. Cysteine-crosslinked oligomeric species were estimated by image scanning and values are presented below the plot in arbitrary units. (C) Effect of NaCl-induced polymerization on DBB vimentin crosslinking. Vimentin wt (3.8 μM final concentration) was incubated with vehicle or 150 mM NaCl for 10 min at 37°C (conditions that induce polymerization), and subsequently incubated for 1 h at r.t. with vehicle or 24 μM DBB. Samples were analyzed by SDS-PAGE and western blot (WB) and representative results are presented. The proportion of oligomeric (bracket) versus monomeric species is shown in the graph at the right. Results are average values of three assays ± SEM. *p<0.05 vs control (−NaCl, −DBB); #p<0.05 vs (−NaCl, +DBB) by paired Student’s *t*-test. (D) Models of vimentin dimers of dimers. Distant-Cys and nearby-Cys tetramers are represented in orange and blue, respectively. For each dimer, 100 ns of MD simulation was performed. The structures of the initial and last frames of the simulation are shown. Cys328 is represented in spheres. (E) Front view of the dimerization interface between both vimentin dimers. (F) Semi-lateral view of both vimentin dimers with details of the interactions that take place at the dimerization interface. Interacting residues are depicted as sticks. (G) Lateral view of the dimerization interface between both vimentin dimers.

In view of the above considerations, we addressed the modeling of a dimer of dimers to study whether the cysteine crosslinking would be geometrically possible (Figure 4D). For this purpose, the vimentin dimer model B, with the cysteine residue in thiol form, was used following protein-protein docking approaches. Several dimer-dimer assemblies were predicted and analyzed, and finally only two tetrameric complexes were selected that accomplished two conditions: 1) they presented an antiparallel orientation, according to the literature ^14, 19^, and 2) the dimer backbones adopted an overall longitudinal alignment. These two “dimers of dimers” differed in the distance between cysteine residues: in the so-called “distant-CYS” arrangement, the cysteine residues were 22 Å apart (Figure 4D, two upper panels), whereas in the “nearby-CYS” arrangement, the distance between cysteine residues was 4 Å (Figure 4D, two lower panels). The stability of these two tetrameric complexes was then studied by running 100 ns MD simulations (Figure S6). The structure of the “distant-CYS” complex, bent during the simulation and the terminal regions of the dimers fell apart (Figure 4D, two upper panels). On the contrary, the “nearby-CYS” tetrameric structure remained stable during the simulation, and only subtle changes in the conformation and distance between dimers were observed (Figure 4D, two lower panels). Therefore, only the “nearby-CYS” arrangement was considered for further calculations, given the fact that the distance between the cysteine residues (around 4 Å) is compatible with the possibility of an oxidative or chemical crosslinking (distance < 6 Å, DBB average crosslinking distance, 4.88 Å, ^70^). Figure 4E through G show additional views of the dimerization interface of the “nearby-CYS” dimer or dimers putatively generated under oxidative conditions. The two antiparallel chains appeared bound through a strong network of ionic interactions, e.g., between the side chains of Arg300 and Glu346*, Lys303 and Glu338*, Arg308-Gln323 and Asp331*, Asp331 and Arg308*-Gln323*, Glu338 and Lys303*, and Glu346 and Arg300* (Figure 4F, G). Nevertheless, our results do not exclude that both, “nearby-Cys” and “distant-Cys” arrangements of the vimentin tetrameric complex and potentially other assemblies, including those proposed from amino group crosslinking studies, coexist (schematized in Figure 1). Vimentin filaments are generally considered to be formed by eight tetramers per section. Early oxidative crosslinking studies of heterooligomers of vimentin and GFAP or desmin proposed that dimers involved in crosslinking would be in a symmetrical “face to face” disposition, implying the establishment of a cysteine disulfide between monomers in the same orientation ^67, 68^. Besides, it was proposed that oxidative crosslinking was always intra-filament. However, later crosslinking studies using amino-reactive reagents have indicated that neighboring dimers within a filament are always antiparallel ^19^, and this has been reflected in several models ^12, 14, 19^ (see scheme in Figure 1). Therefore, several possible arrangements or orientations for the establishment of disulfide-bonded vimentin dimers may exist, although the consideration of all of them falls outside the scope of this study.

### Modeling the interaction of the “nearby-CYS” vimentin tetrameric complex with zinc

Next, the interaction of the putative “nearby-CYS” tetrameric arrangement with zinc ions was studied (Figure 5). To this end, 100 zinc atoms were introduced following the protocol described for the dimer, in order to fill all the possible zinc binding sites, and the systems were submitted to 30 ns MD simulations (Figure S7). A total of 26 zinc atoms remained in contact with vimentin residues more than 90% of the simulation time, confirming the existence of putative zinc hot spots in the vimentin “nearby CYS” tetrameric complex, which in fact double the number estimated for the vimentin dimer. A closer look to the region surrounding Cys328 showed stable interactions of zinc ions with the vicinal Glu329 and Asp331 residues, resembling those observed in the thiol-form of the dimer (Figure 5A).

**Figure 5.**
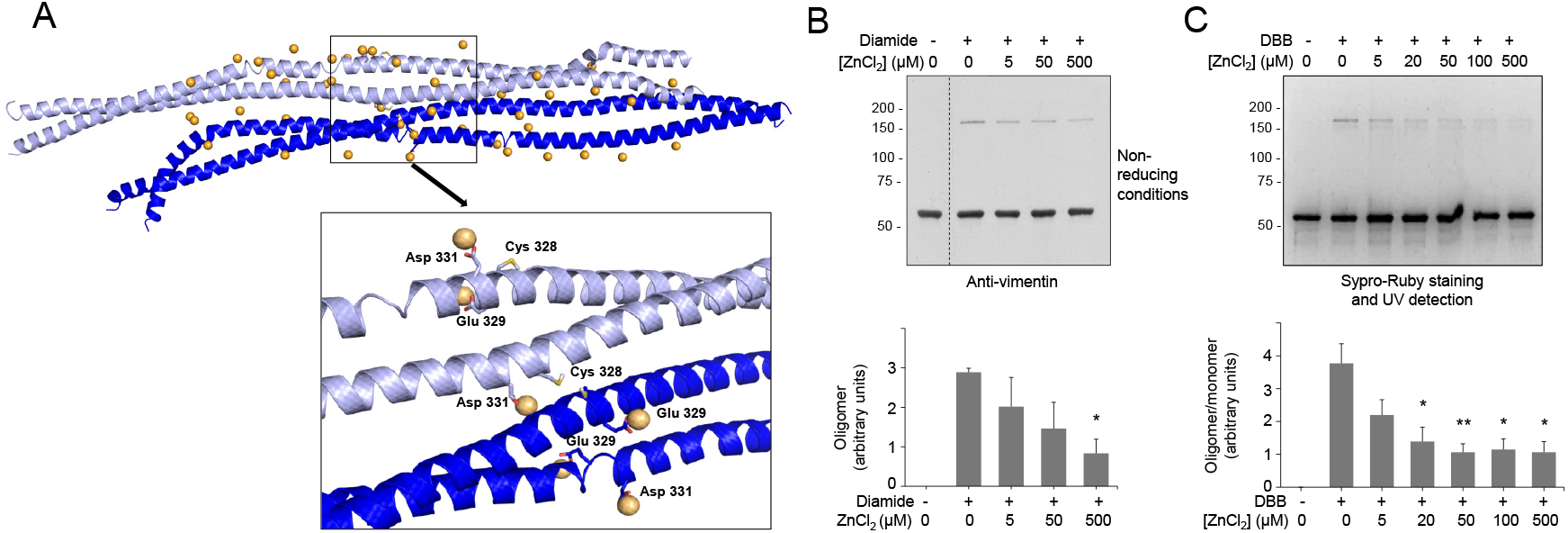
Interaction of zinc with the vimentin nearby-Cys vimentin tetrameric association and effect on cysteine crosslinking. (A) Zinc atoms remaining in the proximity of the vimentin nearby-Cys tetrameric association, with Cys328 in its thiol form are shown after 30 ns of MD simulation. Details of zinc interactions with residues in the vicinity of Cys328 are depicted in the enlarged image. (B,C) Effect of zinc on vimentin cysteine crosslinking. Vimentin wt (4.3 μM final concentration) was incubated in the presence of increasing concentrations of ZnCl_2_ for 1 h at r.t before induction of cysteine crosslinking by oxidation with 1 mM diamide (B) or chemical crosslinking with 24 μM DBB (C), for an additional hour. Incubation mixtures were analyzed by non-reducing SDS-PAGE followed by western blot with anti-vimentin (B) or by SDS-PAGE followed by Sypro-Ruby staining and UV detection (C). The ratios of oligomeric vs monomeric species are depicted in the graphs as average values ± SEM from three assays. *p<0.05 vs diamide or DBB in the absence of ZnCl_2_;**p<0.001 vs DBB in the absence of ZnCl_2_, by unpaired Student’s *t*-test.

Interactions equivalent to those found in the thiolate form of the dimer can also be deduced, as well as for several combinations of cysteine species (thiol or thiolate), which could occur. Therefore, we cannot exclude other dimer-dimer association patterns involving thiol and thiolate species of vimentin simultaneously, in which the coordination of zinc ions will occur through the predominant participation of the carboxylate or the cysteine residues, respectively. In fact, zinc atoms are observed to interact at the interface between both dimers. As more detailed structural information on vimentin filaments becomes available, it will be possible to predict whether zinc atoms participate in protein-protein interactions at the dimer or tetramer levels under physiological conditions, or even in filament bundling, as suggested by recent evidence ^39, 40^.

Next, the Cys328Ser dimer of dimers was constructed as a control. This structure revealed a significant difference with the wt tetrameric assembly consisting in the formation of a H-bond between the OH groups of Ser328 residues from the different dimers, since their side chains fall in the correct orientation and distance (Figure S8). This contrasts with the wt tetrameric assembly, in which the distance between SH groups was longer and oscillated considerably along the MD simulation time. Mutation of the modelled Ser328 back to Cys328 led to a geometry similar to that of the wt, where the distance between SH groups does not correspond to a H-bond distance. Nevertheless, a Cys328Ser vimentin mutant will not undergo cysteine oxidative or chemical crosslinking for which it is not known whether this kind of association would occur.

### Zinc protects vimentin from cysteine-mediated dimerization

Since preincubation with zinc protects vimentin from modification by several electrophilic agents, we explored its effect on cysteine crosslinking. As shown in Figure 5B, diamide-induced disulfide oligomerization of vimentin was diminished by preincubation with ZnCl_2_ in a concentration-dependent manner. Moreover, crosslinking by DBB was markedly attenuated in the presence of zinc (Figure 5C). Importantly, this protective effect could be observed at low micromolar ZnCl_2_ concentrations, that are similar to those of the protein present in the assay, and in the range of the concentrations of zinc found in biological settings, e.g., in plasma ^59^. Interestingly, the protective effect of zinc against cysteine modification and crosslinking affected not only soluble vimentin but also vimentin filaments preformed by addition of NaCl (Figure S9A and B).

### The protective effect of zinc on vimentin is selective

Next, we explored the protective effect of zinc on vimentin crosslinking by other agents. For this purpose, we first used bifunctional cysteine crosslinkers differing in the length of their spacer arms (Figure S10). Interestingly, the protective effect of zinc appeared to inversely correlate with the crosslinkage length resulting in a 76% inhibition of oligomer formation by DBB (4.88 Å), 56% inhibition of crosslinking by tris(2-maleimidoethyl)amine (TMEA, 10.3 Å) and no significant decrease in bismaleimidohexane-induced crosslinking (BMH, 13.0 Å)(Figure 6A).

**Figure 6.**
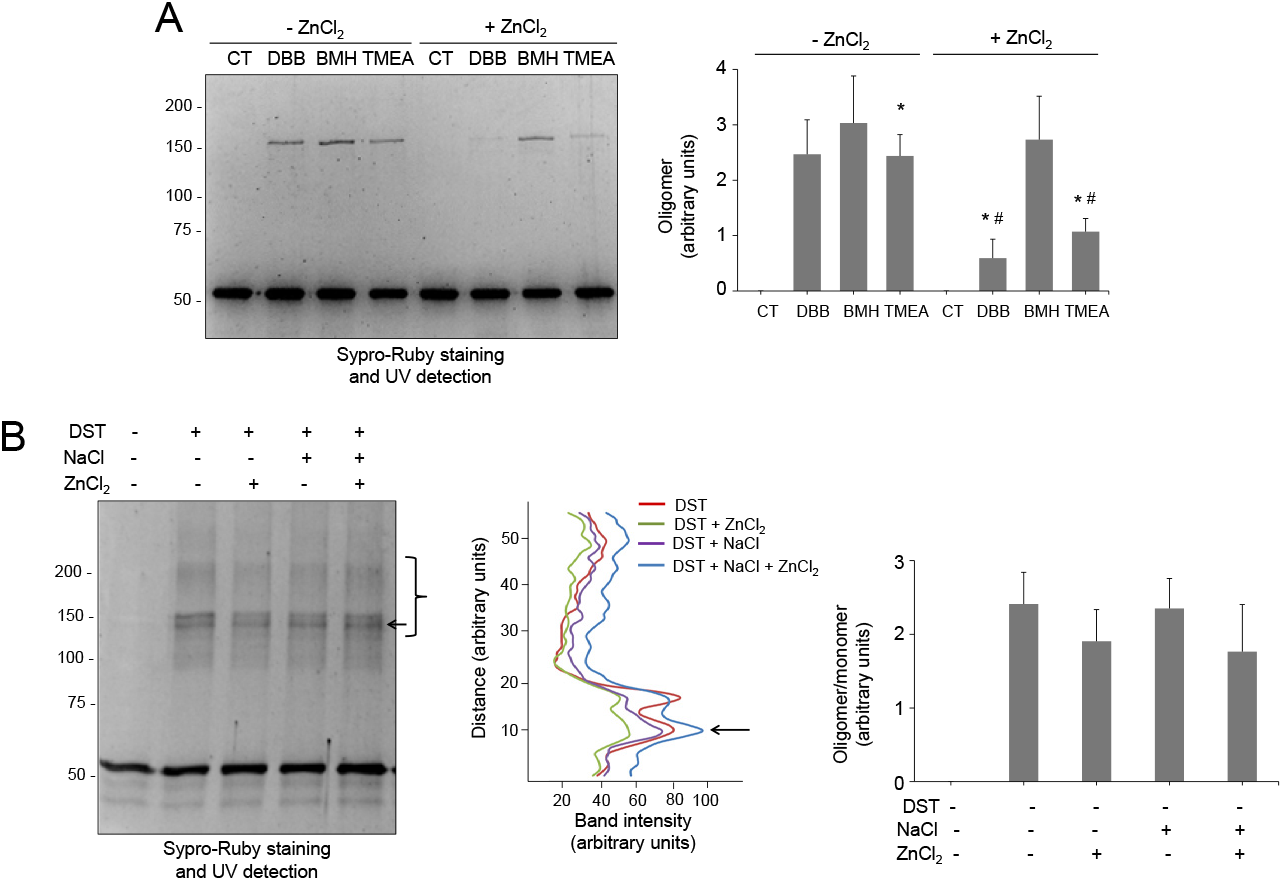
Selectivity of the protective effect of zinc on vimentin crosslinking. (A) Vimentin wt (4.3 μM final concentration) was incubated with vehicle or 500 μM ZnCl_2_ for 1 h at r.t., before addition of vehicle (DMSO) or the indicated cysteine crosslinkers at 24 μM for incubation during 1 h more. Incubation mixtures were analyzed by SDS-PAGE and Sypro-Ruby staining. Oligomeric species were estimated by image scanning and mean values ± SEM from three assays are represented in the graphs. *p<0.05, with respect to the reaction with the same crosslinker in the absence of ZnCl_2_ by paired Student’s t-test. (B) Effect of zinc on vimentin crosslinking by an amino-reactive compound. Left panel, vimentin wt (5.8 μM final concentration) was incubated with vehicle, 150 mM NaCl, or 500 μM ZnCl_2_ for 1 h at r.t. Incubation mixtures were then treated with the amino-reactive crosslinker disuccinimidyl tartrate (DST) at 400 μM for 1 h at 37°C and analyzed as in (A). Middle panel, the intensity of the oligomeric species in the region delimited by the bracket was estimated and is depicted in the graph as a function of the distance (from top to bottom of the marked region). The arrow pointing to the most intense band is shown for reference. Right panel, the ratio of oligomeric (bracket) versus monomeric species was estimated and is depicted in the graph as mean values ± SEM from three assays. No significant differences were found by paired Student’s *t*-test.

These results could suggest that interaction with zinc selectively protects a region of vimentin where cysteine residues from different tetramers or subunits are located at a relatively short distance, i.e., <6 Å, as that measured in the “nearby-CYS” model. Of note, although TMEA is a three arm crosslinker, only vimentin dimers were observed. These results can be interpreted in several ways. First, according to current knowledge, the side chains of cysteine residues in the vimentin coiled-coil dimer would be oriented outwards. Therefore, crosslinking should involve monomers belonging to different dimers or tetramers. In this scenario, the possibility exists that zinc interaction in the proximity of Cys328 sterically precludes DBB crosslinking. Alternatively, zinc interaction with vimentin at this or other sites could induce conformational changes or associations in which cysteine residues are no longer at distances or orientations amenable to crosslinking with DBB or TMEA, while remaining susceptible to BMH-induced crosslinking. Indeed, micromolar zinc induces the association of soluble vimentin into distinct oligomeric assemblies that differ from ULF in their narrower dimensions and in their compromised elongation upon addition of NaCl ^40^. In contrast, when interacting with preformed vimentin filaments, zinc promotes thickening and bundling, which are dependent on the presence of the cysteine residue, without altering filament integrity ^40^. Therefore, these observations indicate that zinc does indeed induce some rearrangement of vimentin structures, at least in part by interacting with Cys328. Nevertheless, the fact that the protective effects of zinc occur both in soluble and filamentous vimentin (Figure S9A and B) suggests that they are not simply due to a tighter packing of the subunits. Moreover, both zinc-induced vimentin oligomerization ^40^ and protection from cysteine modification are reversible upon addition of zinc chelators (Figure S9C), which indicates that the rearrangements do not imply a denaturation of the protein.

As stated above, the products of the crosslinking of vimentin through amino groups have been exhaustively studied, both in soluble vimentin and in filaments. This has led to the definition of the modes of assembly of vimentin tetramers and their position in filaments, schematized in Figure 1A ^19, 71^. In order to get further insight into the specificity of zinc protection, its effect on amino-crosslinking was explored. As previously reported, crosslinking with the bifunctional amino crosslinker disuccinimidyl tartrate (DST, spacer arm, 6.4 Å), induced the appearance of multiple vimentin oligomeric species (Figure 6B, left panel). Interestingly, preincubation with ZnCl_2_ did not diminish DST-induced vimentin crosslinking significantly, nor altered the pattern of oligomeric bands detected, either in the absence or presence of NaCl (Figure 6B, middle panel), suggesting that the protective effect of zinc is selective for thiol crosslinkers of a given length.

### Interaction of vimentin with magnesium

In contrast with the recently explored vimentin-zinc interaction ^4, 72^, the interaction of vimentin with other divalent cations, including Ca^2+^ and Mg^2+^, has been extensively studied ^37, 73^. In particular, magnesium has been reported to induce an electrostatic crosslinking of the vimentin tail segments, which impacts on the physical properties of vimentin filaments ^38, 74^. Therefore, we were interested in assessing the potential binding of magnesium in the area surrounding Cys328. MD simulations of the thiol form of the vimentin dimer in the presence of magnesium ions revealed that, in contrast with the two zinc ions bound in the proximity of the Glu329 and Asp331 hot spot, only one magnesium ion could bind to this region (Figures 7A and S11). This fact could be due to the bigger size of the magnesium ion that would preclude a proper anchorage at the zinc ion site. Indeed, the binding of magnesium to vimentin has been reported to occur mainly at the C-terminal segment of the protein ^37^. Thus, although vimentin behaves as a polyelectrolyte able to bind multiple divalent cations, our results indicate that their mode of binding and effects on the properties of the protein are not identical, with Zn^2+^ exerting distinctive effects in terms of concentration and, potentially, protein conformation ^40^.

**Figure 7.**
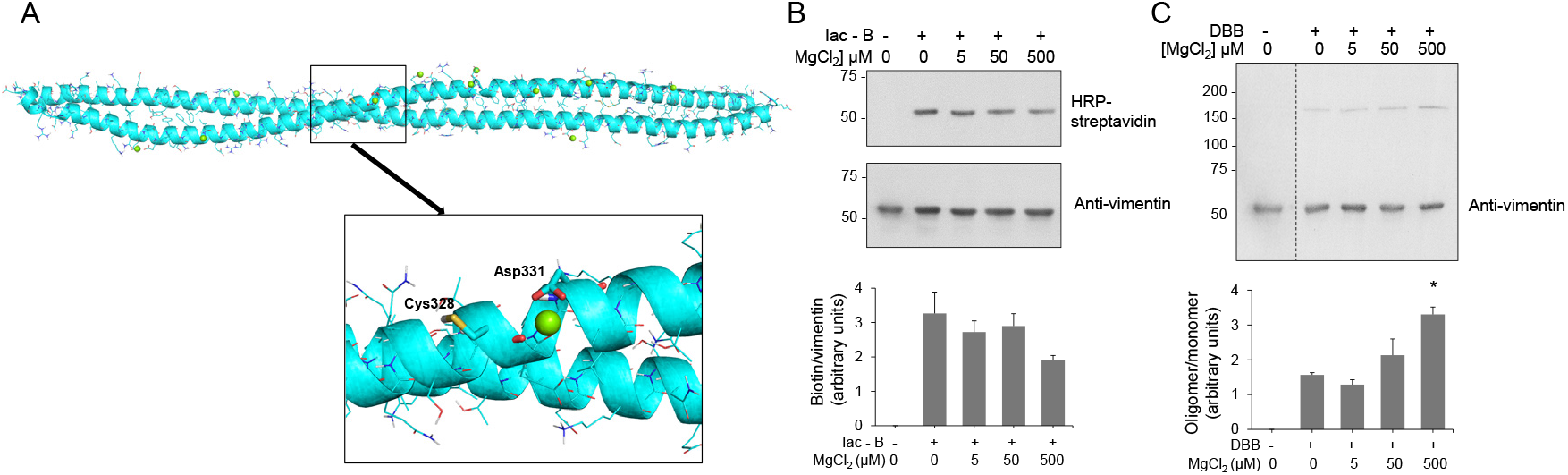
Interaction of magnesium with the vimentin dimer and effect on cysteine crosslinking. (A) Final snapshot of the MD simulation of vimentin dimer B in the presence of magnesium atoms (green spheres). The lower panel shows an enlarged view of the region surrounding Cys328. (B, C) Effect of magnesium on vimentin chemical modification by Iac-B (B) or crosslinking by DBB (C). Vimentin wt (4.3 μM final concentration) was incubated with vehicle or MgCl_2_ at the indicated concentrations for 1 h, at r.t, after which, 10 μM Iac-B (B) or 24 μM DBB (C) were added for 30 min or 1 h, respectively, at r.t. Graphs in the lower panels show mean values ± SEM of at least three independent experiments. *p<0.05 vs DBB by unpaired Student’s *t*-test.

Next, we explored the potential protective effect of magnesium on vimentin cysteine alkylation and crosslinking. Low micromolar concentrations of MgCl_2_ did not diminish the incorporation of Iac-B into vimentin, although higher micromolar concentrations, i.e., 500 μM, tended to decrease Iac-B binding (Figure 7B). Moreover, MgCl_2_ did not exert any protective effect on DBB-induced vimentin crosslinking. Conversely, preincubation with MgCl_2_ at 500 μM (Figure 7C), or even at millimolar concentrations (Figure S12), that is, in a 100-1000-fold molar excess over the protein, increased the intensity of the DBB-induced dimer band. This could be related to the ability of millimolar magnesium to induce vimentin polymerization ^38^, which, as shown in Figure 4C for NaCl-induced polymerization, could facilitate DBB-induced vimentin crosslinking. Moreover, magnesium and calcium are known to mediate lateral interactions favoring filament attraction and aggregation ^75^, and hence they may favor conformations in which Cys328 from different tetramers or filaments become amenable to crosslinking by DBB.

In summary, based on our cysteine modification assays and computational models, we propose that the region surrounding the single cysteine residue of vimentin behaves as a hot spot for zinc binding, possibly modulating vimentin assembly. The proposal of this zinc-binding motif is compatible with the binding of this cation at multiple vimentin sites, and also with the potential contribution of distant segments of the protein, such as the head or tail domains, or of other dimers to zinc binding at the Cys328 region once the protein is organized in filaments. Of note, within the intermediate filament family, keratins have long been known to bind zinc ^76, 77^, although the precise site(s) of interaction, to the best of our knowledge, have not been elucidated. Moreover, Cys328 is not conserved in keratins. Interestingly, other cytoskeletal proteins, including actin and tubulin can bind zinc and are considered zinc-scavenging proteins. In the case of tubulin, zinc can induce various assemblies, including microtubules, sheets and macrotubes, depending on the experimental conditions. Remarkably, zinc appears to induce conformational changes affecting the lateral interactions of tubulin protofilaments, favoring the formation of sheets, in which protofilaments are assembled in an antiparallel fashion ^78^. This associates with zinc-induced conformational changes affecting lateral contacts ^79^ through its interaction with histidine and glutamic residues from adjacent subunits ^80^. Thus, in view of the current evidence, the cysteine hot spot for zinc binding in vimentin and putatively other Type III intermediate filament proteins could represent a selective site of interaction with potential implications in the regulation of these proteins by redox mechanisms.

The pathophysiological consequences of the exposure of vimentin to zinc could be diverse depending on its location. We have previously shown that the vimentin network is thinner and more susceptible to oxidative damage in fibroblasts from patients with genetic zinc deficiency, i.e., acrodermatitis enteropathica ^81^, and these cellular alterations are ameliorated by zinc supplementation ^4^. In addition, zinc is able to induce aggregation of vimentin and other proteins ^40, 82^, which could contribute to pathologies associated with extracellular accumulations of vimentin and zinc, as the pathological deposits that develop in diseases such as age-related macular degeneration ^83^. Moreover, in a cellular context, zinc availability will influence vimentin dynamics by multiple mechanisms, including the control of enzymes modulating cellular redox status, zinc-dependent proteases or transcription factors ^44^.

### Concluding remarks

Compared to the other main cytoskeletal systems, microfilaments and microtubules, the regulation of the assembly and reorganization of intermediate filaments is insufficiently understood. Apart from posttranslational modifications, interaction of intermediate filaments with “cofactors”, and specifically with divalent cations, is arising as a potential regulatory mechanism in vitro and in cells. In previous works, we identified the interaction of micromolar concentrations of zinc with vimentin and its impact on vimentin network organization and filament morphology, in vitro as well as in cellular models of disease. In addition, zinc, but not other divalent cations, affected the response of vimentin to oxidants and electrophiles, for which the presence of its single cysteine residue is an important factor. The findings herein reported provide a biochemical basis for the behavior of vimentin and, potentially, other members of the Type-III intermediate filament family, as redox sensors relying on the particular characteristics of the cysteine residue. Moreover, the identification of a hot spot for zinc binding in the region of the cysteine residue supports the role of zinc in vimentin organization and redox regulation, and offers a working hypothesis to assess the role Type-III intermediate filaments in the pathogenesis of diseases associated with oxidative or electrophilic stress and/or zinc deficiency.

## Materials and Methods

### Materials

Dimethylsulfoxide (DMSO), monosodium and disodium phosphates, Tris, Pipes, DTT, PMSF, diamide and dibromobimane (DBB, spacer arm, 4.88 Å ^70^) were from Sigma. Tris(2-maelimidoethyl)amine, (TMEA, spacer arm, 10.3 Å) and bismaleimidohexane (BMH, spacer arm, 13.0 Å) were from Thermo Fisher Scientific. Monobromobimane (MBB, sc-214629) and (+)-Biotin-(PEO)3-iodoacetamide (biotinylated iodoacetamide, Iac-B) were purchased from Santa Cruz Biotechnology. Citric acid, sodium carbonate and bicarbonate, 2-mercaptoethanol and EDTA were obtained from Merck. Precision Plus-Dual Color protein standards and Sypro Ruby were from BioRad. The biotinylated analog of 15-deoxy-Δ^12,14^-prostaglandin J_2_ (15d-PGJ_2_-B) was from Cayman Chemical. No unexpected or unusually high safety hazards were encountered throughout this work.

### Computational Methods

#### Building of the vimentin models

PyMol software ^84^ was used to build the 3D models from the vimentin X-ray structures available at the Protein Data Bank ^85^ (https://www.rcsb.org, last access October 14th), with PDB IDs 3KLT and 1GK4. The sequences of PDBs 3KLT and 1GK4 structures share seven amino acids (Cys328, Glu329, Val330, Asp331, Ala332, Leu333 and Lys334). One monomer of each PDB structure was kept and superimposed to the dimer from the other PDB structure by this overlapping region. Thus, two vimentin dimer models comprising residues Asp264 through Gly406, but differing in their geometry, were constructed: 1GK4 superimposed to 3KLT model (model A) and 3KLT superimposed to 1GK4 model (model B). Only model B was considered for further computational studies.

### Protein-protein docking

Model B of vimentin dimer from the molecular dynamics (MD) simulations was submitted to servers HADDOCK ^86^ and ZDOCK ^87^ to perform protein-protein docking. A total of 20 vimentin dimers of dimers were obtained and analyzed.

### Molecular Dynamics (MD) simulations

MD simulations were carried out using Amber14. Eight steps of preparation were performed before running MDs. The first one consisted of 1000 steps of steepest descent algorithm followed by 7000 steps of conjugate gradient algorithm; a 100 kcal·mol^−1^·A^−2^ harmonic potential constraint was applied on both the proteins and the ligand. In the four subsequent steps, the harmonic potential was progressively lowered respectively to 10, 5, and 2.5 kcal·mol^−1^·A^−2^ for 600 steps of conjugate gradient algorithm each time, and then the whole system was minimized uniformly. In the following step the system was heated from 0 to 100 K using the Langevin thermostat in the canonical ensemble (NVT) while applying a 20 kcal· mol^−1^·A^−2^ harmonic potential restraint on the proteins and the ligand. The next step heated up the system from 100 to 300 K in the isothermal−isobaric ensemble (NPT) under the same restraint condition as in the previous step. In the last step, the same parameters were used to simulate the system for 100 ps, but no harmonic restraint was applied. At this point the system was ready for the production run, which was performed using the Langevin thermostat under NPT ensemble, at a 2 fs time step. All production runs were performed for 100 or 150 ns. Zinc atoms were added to the system using leap.

### Computational estimation of Cys328 pKa

The models of vimentin dimer B and the corresponding vimentin dimer of dimers assembly were used for calculations. PROPKA ^88, 89^ was employed to predict te pKa of Cys328 at physiological pH (pH=7).

### Vimentin refolding

Recombinant human vimentin wild type (wt) and Cys328Ser in 8 M urea, 5 mM Tris-HCl pH 7.6, 1 mM EDTA, 10 mM 2‑mercaptoethanol, 0.4 mM PMSF and approximately 0.2 M KCl, purified essentially as described ^90^, were purchased from Biomedal (Spain). The proteins were ultrafiltrated using 10 K pore size Amicon Ultra filter units (Millipore) and step-wise dialyzed against 5 mM Pipes-Na pH 7.0, 1 mM DTT containing decreasing urea concentrations (8, 6 and 2 M). Final dialysis was performed for 16 h at 16°C in 5 mM Pipes-Na, pH 7.0, 0.25 mM DTT ^91^. Ultrafiltration was necessary for efficient removal of EDTA ^91^. Protein concentration was estimated from its A_280_ nm, using an extinction coefficient of 22450 M^−1^cm^−1^. Aliquots of the protein were kept at −80°C until use.

### Experimental determination of the cysteine pKa

Buffers were prepared by mixing 5 mM concentrations of each of the following to obtain the desired pH: Citric acid/disodium phosphate (pH 3-5.5); disodium phosphate/monosodium phosphate (pH 6-8); Tris/HCl (pH 8.5-9); sodium carbonate/sodium bicarbonate (pH 9.5-10.8). Purified vimentin wt or Cys328Ser were thawed and subjected to five cycles of dilution with 5 mM Pipes/Na pH 7.0 (10-12 ml) and ultrafiltration on Amicon Ultra-15 units (10 K cutoff; Millipore) for DTT elimination, immediately before monobromobimane (MBB) modification. The last flowthrough was kept and the protein concentration was measured on a Nanodrop (Thermo Fisher). MBB 50 mM was prepared fresh in DMSO and later diluted with 5 mM Pipes/Na pH 7.0 to 333 μM, the concentration required to obtain a final 1:4 (mol/mol) ratio protein:MBB in the modification protocol. MBB reactions were carried out in triplicate in the dark on black-polystyrene NBS-treated multiwell plates (ref. 3650, Corning) in a final volume of 100 μl/well. Each well contained 2.5 μM vimentin wt or Cys328Ser, 77 μl of buffer at the desired pH and 10 μM MBB. Controls lacking the protein included the flowthrough of ultrafiltration to establish the background or reduced glutathione (G4251, Sigma) as positive control of MBB modification. Kinetics of modification were followed by measuring fluorescence emission at 480 nm every 3 min for up to 90 min, upon excitation at 380 nm using Varioskan Flash (Thermo Fisher). The final pH for each reaction was measured with a pH-meter using equivalent mixtures in a final volume of 2 ml. Fluorescence data for each well were adjusted to a sigmoidal curve using GraphPad Prism v.5 to obtain the value of the bottom plateau corresponding to the initial non-alkylated protein (F_o_). The ratio of alkylated protein at time t (F_t_) vs F_o_ was calculated and used to obtain the ln(F_t_/F_o_). The slopes of the representation of ln(F_t_/F_o_) against time provide the k_obs_ values, which were represented against the final reaction pH to obtain the pKa (the inflexion point of the curve). Three independent experiments were carried out for both vimentin wt and Cys328Ser using different protein preparations.

Additional verification of the results obtained with MBB was achieved by subjecting vimentin to modification with Iac-B at different pHs for 30 min at room temperature, in the dark. Reactions contained 2.5 μM vimentin wt or Cys328Ser, 50 μM Iac-B and buffers at different pHs near the pKa values calculated in the MBB modification assays. Reactions were stopped by boiling in Laemmli sample buffer. Proteins were separated in 10% SDS-polyacrylamide gels and electrotransferred to PVDF membranes using a semi-dry system (Bio-Rad). Biotin incorporation was assessed by incubating blots with streptavidin-HRP (1:1000 v/v, GE Healthcare), and vimentin levels estimated by western blot with anti-vimentin V9 monoclonal antibody (1:1000 v/v; sc-6260, Santa Cruz), followed by enhanced chemiluminescent (ECL) detection (GE Healthcare). Signal intensities were evaluated by image scanning and analysis with ImageJ.

### Vimentin modification by oxidants, electrophiles and crosslinkers

Modification of vimentin was assessed by gel-based techniques. Briefly, vimentin at 3.8 μM in 5 mM Pipes-Na, pH 7.0 was incubated for 1 h at room temperature in the presence of the indicated compounds. Given the fact that DTT can react with electrophilic compounds and chelate metals ^92^ its final concentration was kept below 0.2 mM. To evaluate the protective effect of the different salts on vimentin modification or crosslinking, the protein was preincubated with ZnCl_2_, MgCl_2_ or NaCl, for 1 h at room temperature, as indicated. Incorporation of 15d-PGJ_2_-B or Iac-B was analyzed by SDS-PAGE followed by blot and biotin detection as described above. For detection of crosslinked products, samples were run on 7.5% SDS-polyacrylamide gels, which were fixed with 40% (v/v) methanol, 10% (v/v) acetic acid for 60 min, stained with Sypro Ruby overnight, washed with 10% (v/v) methanol, 6% (v/v) acetic acid, and visualized under UV light on a Gel-Doc XR Imaging System (Bio-Rad).

### Statistical analysis

All experiments were performed at least three times using different protein batches. Results are presented as average values ± standard error of mean (SEM). Statistical analyses were carried out using GraphPad Prism v5. For comparisons of different sets of values the Student’s *t*-test was used. Differences were considered significant for p<0.05.

## Acknowledgements

We are indebted to MJ Carrasco for valuable technical assistance. Feedback from COST Action CA15214 “EuroCellNet” is gratefully acknowledged.

## Funding

This work was supported by the European Union’s Horizon 2020 research and innovation program under the Marie Sklodowska-Curie Grant Agreement No. 675132 “Masstrplan”, Grants SAF2015-68590-R and RTI2018-097624-B-I00 from Agencia Estatal de Investigación, MCI/FEDER, Spain, and Instituto de Salud Carlos III/FEDER, RETIC Aradyal RD16/0006/0021 to DPS; Grant CTQ2017-88353-R to SMS; Grant BES-2015-071588 to JGC.

## Notes

### Competing Interest Statement

The authors have declared no competing interest.

