## Supporting Information for "Elucidating vimentin interaction with zinc ions and its interplay with oxidative modifications through crosslinking assays and molecular dynamics simulations"

Table of Contents

**Figure S1.** MD simulations of vimentin dimer A (100 ns) and vimentin dimer B (200 ns). RMSD plots.

**Figure S2.** MD simulation of vimentin dimer B with Zinc ions (30 ns). RMSD plot.

**Figure S3.** MD simulation of vimentin dimer B with Cys328 thiol group as thiolate and in presence of Zinc ions (50 ns). RMSD plot.

**Figure S4.** Determination of Cys328 pKa through various methods.

**Figure S5.** MD simulation of vimentin dimer Cys328Ser mutant (50 ns). RMSD plot.

**Figure S6.** MD simulations of vimentin tetrameric complexes (100 ns). A) "distant-CYS" (Cys residues at a distance of 22 Å). B) "nearby-CYS" (Cys residues at a distance of 4 Å). RMSD plots.

**Figure S7.** MD simulation of vimentin tetrameric "nearby-CYS" complex with Zinc ions (100 ns). RMSD plot.

**Figure S8.** MD simulation of vimentin tetrameric Cys328Ser mutant complex (100 ns). RMSD plot.

**Figure S9.** Protective effect of zinc on Cys328 modification and crosslinking in soluble and polymerized vimentin

**Figure S10.** Cysteine crosslinkers used in this study.

**Figure S11.** MD simulation of vimentin dimer B with Magnesium ions (100 ns). RMSD plot.

**Figure S12.** Effect of millimolar magnesium concentration on the crosslinking of vimentin by DBB.

### Supplementary Figures

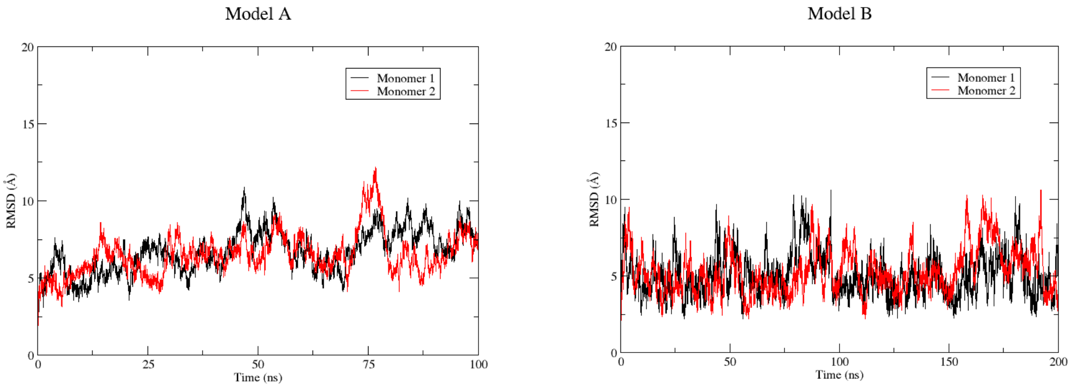


**Figure S1.** MD simulations of vimentin dimer A (100 ns) and vimentin dimer B (200 ns). RMSD plots for the α-Carbons.

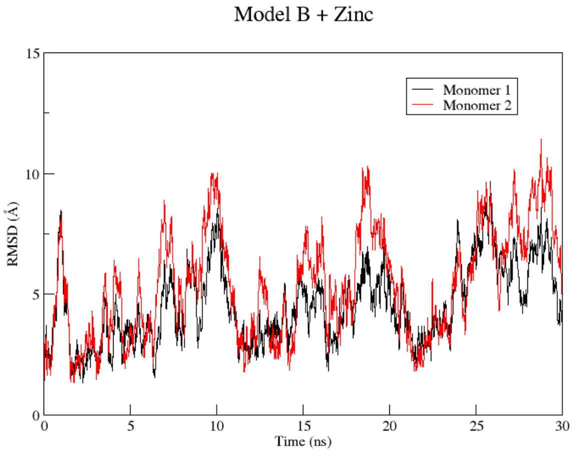


**Figure S2.** MD simulation of vimentin dimer B with Zinc ions (30 ns). RMSD plot for the α-Carbons.

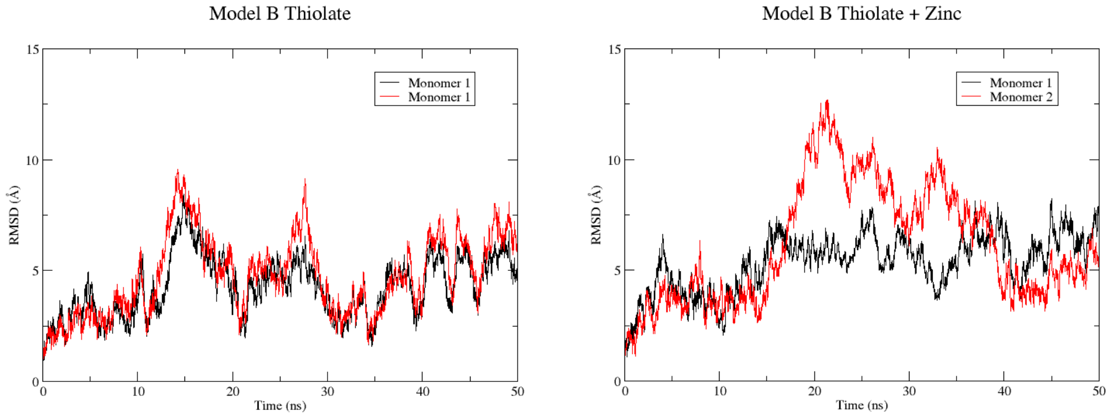


**Figure S3.** MD simulation of vimentin dimer B with Cys328 thiol group as thiolate and in presence of Zinc ions (50 ns). RMSD plot for the α-Carbons.

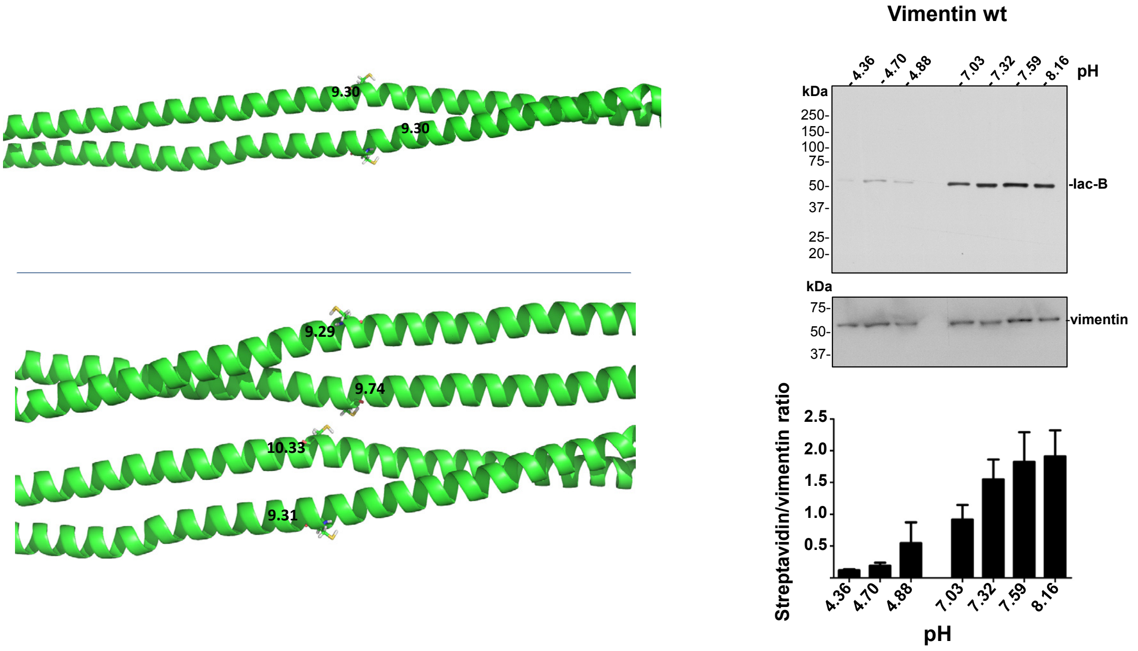


**Figure S4.** Determination of Cys328 pKa through various methods. Computational methods were used to calculate C328S, as detailed in Methods in the context of the vimentin dimer or Nearby CYS tetrameric complex (left panels). Right panel, experimental determination of Cys328Ser pKa through the incorporation of biotinylated iodoacetamide. Western blot analysis of vimentin wt (0.8 μg) modification with biotinylated-iodoacetamide (Iac-B) at pH ranges around the pKa values measured with monobromobimane (please see main text). Biotin incorporation was estimated using streptavidin-HRP (upper blot) and vimentin levels were estimated by western blot with anti-vimentin V9 antibody (lower blot). Densitometric scanning was performed with ImageJ and the streptavidin/vimentin signal ratio obtained as shown in the histogram. The figure shows the mean ± SEM of three independent experiments.

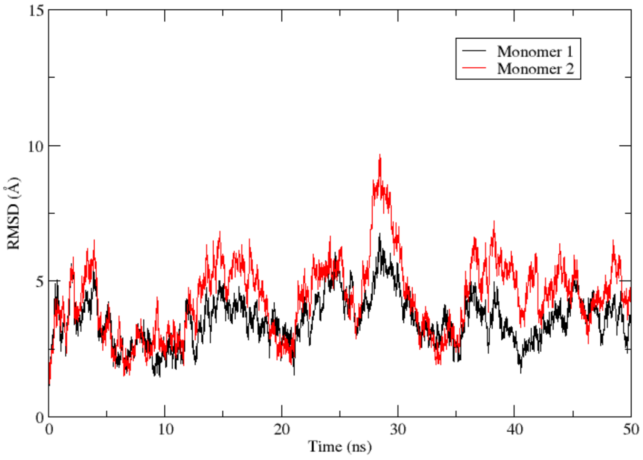


**Figure S5.** MD simulation of vimentin dimer Cys328Ser mutant (50 ns). RMSD plot for the α-Carbons.

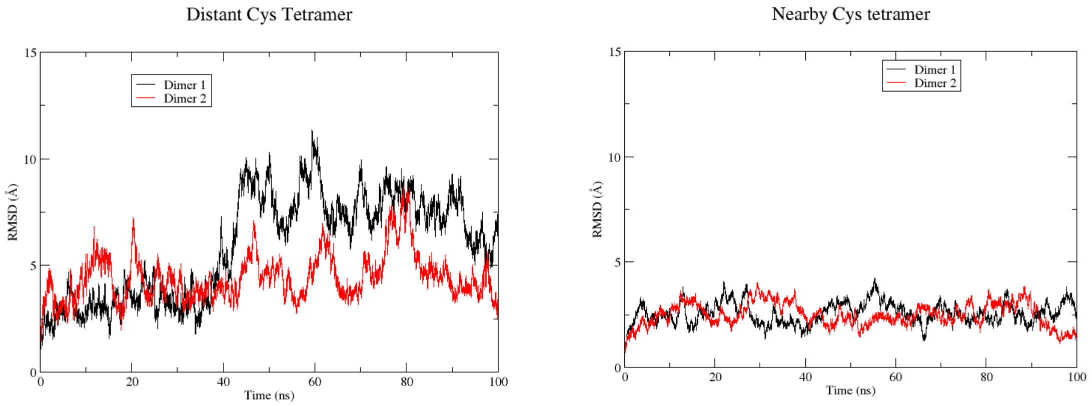


**Figure S6.** MD simulations of vimentin tetrameric complexes (100 ns). A) “distant-CYS” (Cys residues at a distance of 22 Å). B) “nearby-CYS” (Cys residues at a distance of 4 Å). RMSD plots for the α-Carbons.

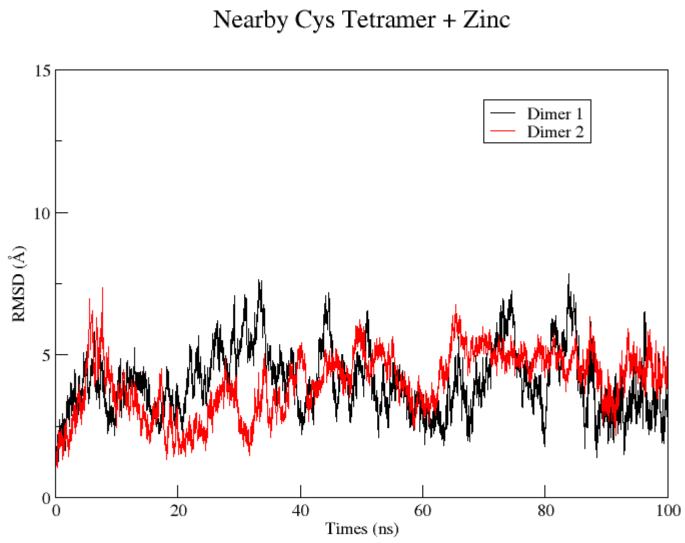


**Figure S7.** MD simulation of vimentin tetrameric "nearby-CYS" complex with Zinc ions (100 ns). RMSD plot for the α-Carbons.

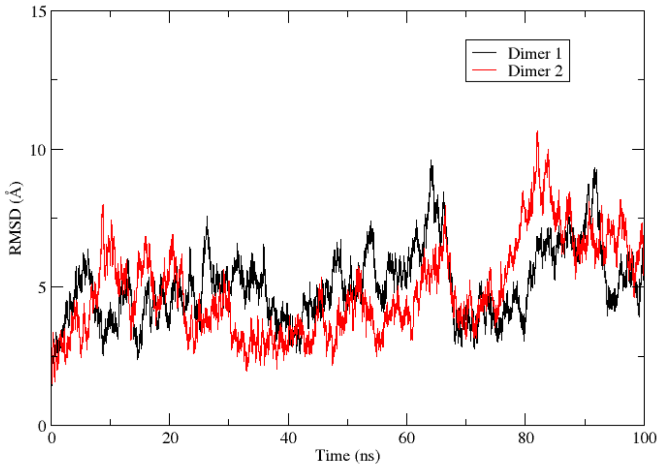


**Figure S8.** MD simulation of vimentin tetrameric Cys328Ser mutant complex (100 ns). RMSD plot.

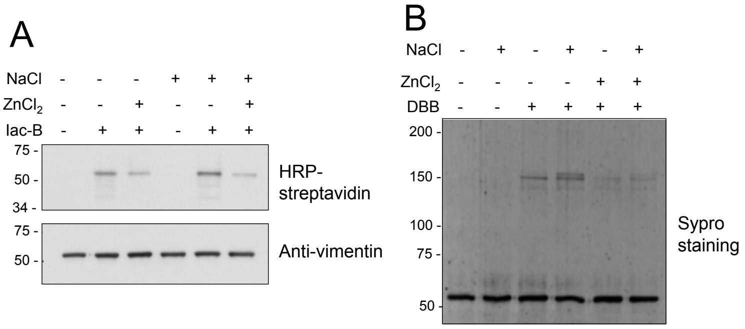


**Figure S9.** Protective effect of zinc on Cys328 modification and crosslinking in soluble and polymerized vimentin. (A) Vimentin at 5 μM was incubated with vehicle or 150 mM NaCl for 10 min at 37ºC, followed by an 1 h incubation with 500 μM $ZnCl_2$, after which, 10 μM biotinylated iodoacetamide (Iac-B) was added for an additional hour at 37ºC. Samples were analyzed by SDS-PAGE and electroblotted. Incorporation of biotin was assessed by detection with HRP-streptavidin and vimentin levels by western blot. (B) Vimentin was incubated in the absence or presence of NaCl followed by $ZnCl_2$ as in (A) and subsequently treated with vehicle or 24 μM DBB for 1 h t r.t. Samples were analyzed by SDS-PAGE and Sypro-Ruby staining. (C) Vimentin was incubated in the presence or absence of $ZnCl_2$, after which the zinc chelator N,N,N′,N′-tetrakis(2-pyridylmethyl)ethylenediamine (TPEN) was added at 1 mM for 30 min before treatment with Iac-B. Incorporation of biotin was detected as in (A). Results are representative from (A) tree, (B) two and (C) two assays.

| Compound | Structure | Spacer (Å) |
|---|---|---|
| Dibromobimane (DBB) | 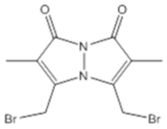 | 4.9 |
| Tris(2-maleimidoethyl)amine (TMEA) | 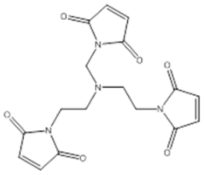 | 10.3 |
| Bismaleimidohexane (BMH) | 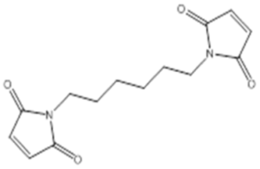 | 13.0 |

**Figure S10.** Cysteine crosslinkers used in this study. Structures of three cysteine-crosslinkers, dibromobimane (DBB), tris(2-maleimidoethyl)amine (TMEA) and bismaleimidohexane (BMH), with the corresponding space arm lengths as specified by the manufacturer.

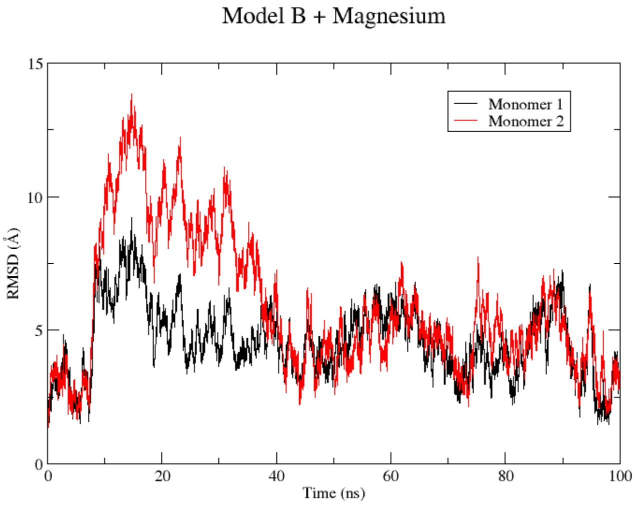


**Figure S11.** MD simulation of vimentin dimer B with Magnesium ions (100 ns). RMSD plot for the α-Carbons.

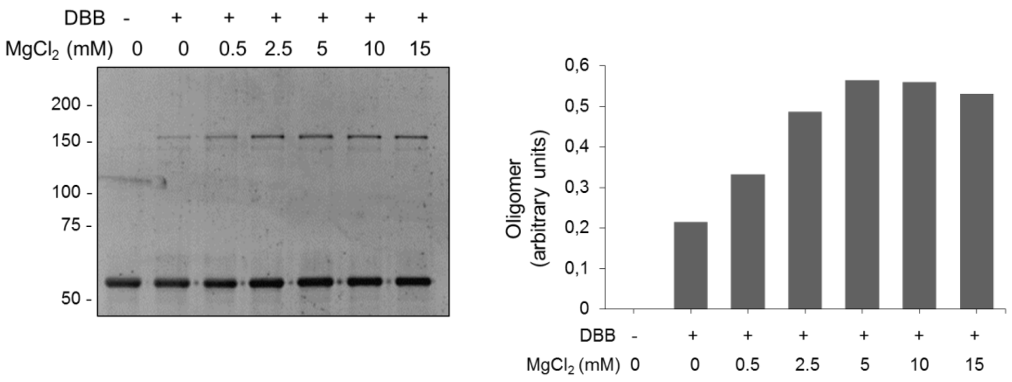


**Figure S12.** Effect of millimolar magnesium concentration on the crosslinking of vimentin by DBB. Vimentin wt at 4.3 μM was first incubated with the indicated concentrations of $MgCl_2$ for 1 h, at r.t, after which, it was incubated with DBB at 24 μM for 1 h more. Incubation mixtures were analysed by SDS-PAGE and protein species visualized by under UV light after SYPRO Ruby staining. Levels of oligomer were estimated by image scanning and are shown in the histogram at the right.
